# Human Ribosomal G-Quadruplexes Regulate Heme Bioavailability

**DOI:** 10.1101/2020.04.15.042721

**Authors:** Santi Mestre-Fos, Chieri Ito, Courtney M. Moore, Amit R. Reddi, Loren Dean Williams

## Abstract

The *in vitro* formation of stable G-quadruplexes (G4s) in human ribosomal RNA (rRNA) was recently reported. However, their formation in cells and their cellular roles have not been resolved. Here, by taking a chemical biology approach that integrates results from immunofluorescence, G4 ligands, heme affinity reagents, and a genetically encoded fluorescent heme sensor, we report that human ribosomes can form G4s *in vivo* that regulate heme bioavailability. Immunofluorescence experiments indicate that the vast majority of extra-nuclear G4s are associated with rRNA. Moreover, titrating human cells with a G4 ligand alters the ability of ribosomes to bind heme and disrupts cellular heme bioavailability as measured by a genetically encoded fluorescent heme sensor. Overall, these results suggest ribosomes are central hubs of heme metabolism.

## INTRODUCTION

Cells tightly control heme concentration and bioavailability (1-3) because it is essential but potentially cytotoxic. Proteins that regulate heme concentration are relatively well understood; structures and mechanisms of all eight heme biosynthetic enzymes and the heme degrading heme oxygenases are known (1-3). However, regulation of heme bioavailability, including intracellular trafficking from sites of synthesis in the mitochondrial matrix or uptake at the plasma membrane, is poorly understood. Current paradigms for heme trafficking and mobilization involves heme transfer by unknown proteinaceous factors and largely ignore contributions from nucleic acids. Given that the first opportunity for protein hemylation occurs during or just after translation, ribosomal RNA (rRNA) or proteins (rProteins) may be critical for shepherding labile heme to newly synthesized proteins.

We hypothesized that intracellular heme bioavailability is regulated in part by rRNA quadruplexes (G4s). G4s are nucleic acid secondary structures that are composed of four guanine columns surrounding a central cavity that sequesters monovalent cations. Our hypothesis is based on the high affinity of heme for G4s (*K*_D_ ∼10 nM) (4-6), our work demonstrating that rRNA forms extensive G-tracts *in vitro* (7, 8), the extreme stabilities of rRNA G4s *in vitro* (7, 8) and the extraordinary abundance of rRNA *in vivo* (9).

DNA G4s are proposed to help regulate replication (10), transcription (11), and genomic stability (12). In RNA, G4s are associated with untranslated regions of mRNA and have been proposed to regulate translation (13-15). However, the *in vivo* folding state and functional roles of G4s are under debate. Eukaryotic cells contain helicases that appear to unfold RNA G4s (16) although counter arguments have been put forth (17, 18). The density of G4 sequences on surfaces of the human ribosome, which is extremely abundant, is high, with 17 G4 sequences in the 28S rRNA and 3 in 18S rRNA (**Figure 1A**). Previous to this report, it was not known if human ribosomes form G4s *in vivo* or what their functions might be.

**Figure 1.**
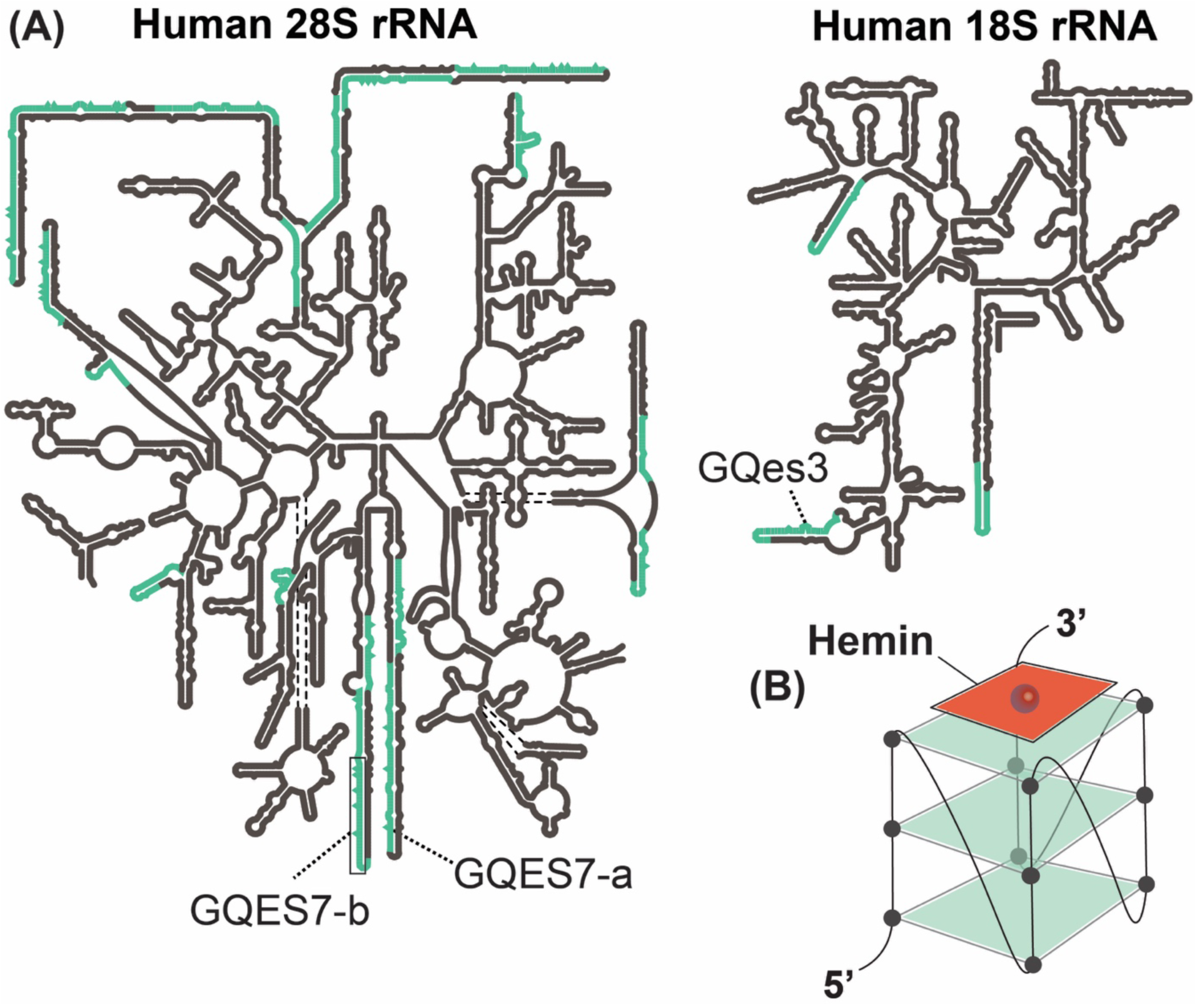
(A) Secondary structures of the human LSU rRNAs (5.8S and 28S) and SSU rRNA (18S). G4 sequences are highlighted in green. rRNA-based oligomers from the LSU (GQES7-a, GQES7-b) and from the SSU (GQes3) are indicated. (B) Schematic representation of a hemin-G4 complex.

Indeed, herein we present evidence that rRNA forms G4s *in vivo* that regulate cellular heme homeostasis. Results of immunofluorescence experiments with a G4 antibody, RNA pulldowns and competition experiments with G4 ligands provide strong support for *in vivo* formation of G4s by rRNA tentacles. We find that G4s on ribosomes bind heme *in vitro* (**Figure 1B**) and that perturbation of G4s *in vivo* with G4 ligands affects *in vivo* heme interactions and heme bioavailability, as measured by heme affinity reagents and genetically encoded heme sensors. Taken together, the results here indicate that surface-exposed rRNA G4s interact with heme in cells and suggest that ribosomes are hubs for cellular heme metabolism.

## RESULTS

### Ribosomal RNA forms G4s in vivo

Confocal microscopy and G4-pulldowns were used to determine if human ribosomes form G4s *in vivo*. For confocal microscopy, we used the BG4 antibody, which selectively targets G4s (19, 20) and has been broadly used for visualizing DNA G4s and non-ribosomal RNA G4s in cells. (20-23) Our method of permeabilizing cells for antibody treatment does not permeabilize the nuclei (24). Therefore, DNA G4s were not anticipated or observed. To identify ribosome associated G4s, we determined the extent to which antibodies to rProtein L19 (eL19) and to G4s colocalize and how this is altered when cells are subjected to RNase or G4 ligand PhenDC3, which are expected to modulate G4-L19 colocalization. Prior to antibody addition, cells were crosslinked with paraformaldehyde to lock G4s *in situ*. This procedure is intended to prevent induction of G4s by the antibody and has been shown to reduce levels of detection of G4s (18). The extent of L19 and G4 antibody colocalization suggests that a fraction of ribosomes form G4s (**Figure 2A,C**) and that most G4s are associated with ribosomes. Specifically, we find that ∼83% of BG4 pixels colocalize with L19, indicating that the vast majority of G4s *in vivo* are associated with ribosomes (**Figure 2C**, green bar) and are therefore rRNA G4s. Conversely, only 5% of L19 pixels colocalize with BG4 (**Figure 2C**, WT red bar), indicating that only a specialized fraction of ribosomes contains G4s. Similar results were obtained using an antibody against rProtein uL4 instead of L19 (not shown).

**Figure 2.**
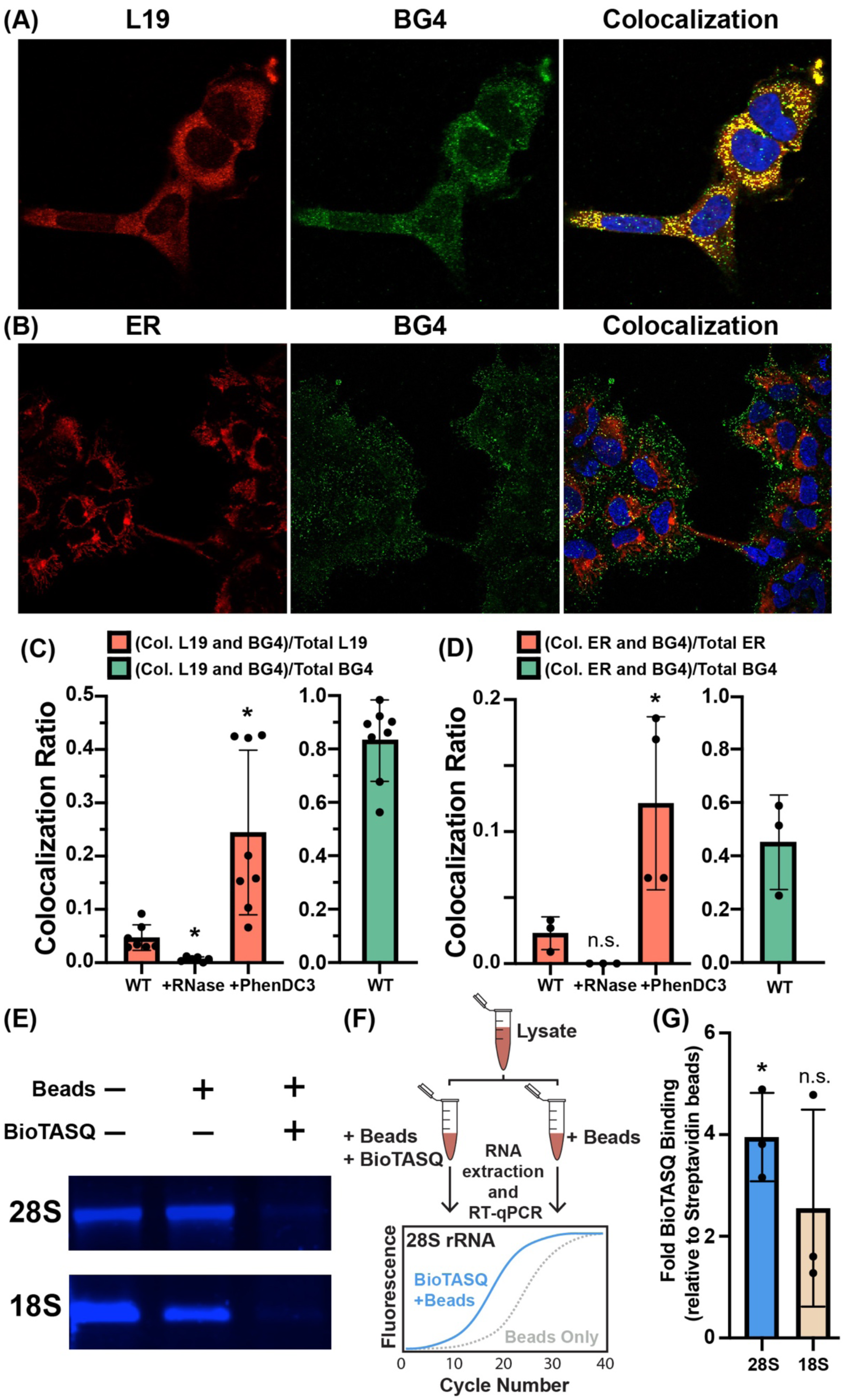
Ribosomal G4s in HEK293 cells. Colocalization of (A) ribosomal protein L19 or (B) endoplasmic reticulum (red) with RNA G4s (green). Nuclei were stained with DAPI (blue). (C) Extent of colocalization is quantitated as the ratio of colocalized pixels over total L19 pixels (red bars) or as the ratio of colocalized pixels over total BG4 pixels (green bar). Same analysis was done for ER-BG4 colocalization (D). The statistical significance relative to WT is indicated by asterisks using an ordinary one-way ANOVA with Dunnett’s post-hoc test. Each dot represents a biological replicate. (E) G4 ligand BioTASQ binds to 28S and 18S rRNAs. In the presence of BioTASQ and streptavidin beads, human rRNAs do not enter the agarose gel. (F) Schematic representation of the BioTASQ pulldown protocol. (G) RT-qPCR analysis of rRNAs pulled down by BioTASQ. The statistical significance relative to a fold enrichment value of 1 is indicated by asterisks using a one sample *t* and Wilcoxon test. Each dot represents a biological replicate. Data in (G) are represented as RNA enrichment under “BioTASQ + streptavidin beads” conditions relative to control streptavidin beads. * P < 0.05. n.s. = not significant.

PhenDC3, which is known to induce and stabilize G4s, (25, 26) appears to increase ribosomal G4 formation *in vivo*; treating cells with PhenDC3 increases L19-BG4 colocalization from 5 to ∼24% (**Figure 2C**). The increase in colocalization upon PhenDC3 treatment supports formation of G4s by ribosomes. By contrast, treating cells with RNase A abolishes the L19-BG4 colocalization signal (**Figure 2C**). Together, these results indicate the colocalized BG4 signal is coming from a G4 forming RNA in close proximity to L19.

mRNA in the cytosol, in the unlikely event that they form G4s at high frequency (16), may confound our ability to selectively detect rRNA G4s. The high density of ribosomes on the surface of the endoplasmic reticulum (ER) and the lower abundance of mRNA in this location as compared to the cytosol (27) motivated us to investigate if G4s colocalize with the ER. Toward this end, we determined the extent to which BG4 colocalizes with an antibody against an ER membrane protein (calnexin) (**Figure 2B**). Indeed, we find that ∼45% of the BG4 signal colocalizes with the ER marker (**Figure 2D**, green bar), indicating a significant presence of RNA G4s at the ER membrane. As with L19, the fraction of the ER signal that colocalizes with G4s (∼2%) is completely abolished by RNase (undetectable) and enhanced by PhenDC3 (12%) (**Figure 2D**). Altogether, the data are consistent with formation of RNA G4s by ER-bound ribosomes.

In an orthogonal approach, we pulled down RNA with BioTASQ (18, 28), which is a G4 ligand linked to biotin. BioTASQ captures G4s. We previously used BioTASQ to demonstrate that human rRNA forms G4s *in vitro* (**Figure 2E**) (8). Here, we captured rRNA G4s from crosslinked HEK293 cells by methods summarized in **Figure 2F**. BioTASQ captures 28S rRNA from cell lysates (**Figure 2G**), in agreement with our previous *in vitro* BioTASQ data and with observations of G4-L19 colocalization above. BioTASQ also captures 18S rRNA although the signal is significantly weaker. Taken together, our immunofluorescence and BioTASQ pulldown experiments provide strong evidence that human ribosomes form G4s *in vivo*.

### Human ribosomes bind hemin in vitro

It has been suggested that G4s might associate with heme *in vivo* (29). *In vitro*, heme binds with high affinity to G4s by end-stacking (30-32) (**Figure 1B**). We used UV-visible spectroscopy to assay the binding of hemin to human rRNA. rRNA oligomers GQES7-a (**Figure 3A**), GQES7-b (**Figure S.1A**) or GQes3 (**Figure S.1B**) were titrated into fixed amount of hemin. GQES7-a and GQES7-b are fragments of expansion segment 7 of human LSU rRNA (7). GQes3 is a fragment of expansion segment 3 of human SSU rRNA (8). Each of these oligonucleotides is known to form G4s and each caused a pronounced increase in the Soret band of hemin at 400 nm. The binding is specific for G4s as a mutant oligonucleotide, *mut*es3, that lacks G-tracts does not induce a change in the hemin Soret band (**Figure S.1C**). Larger human ribosomal components also bind heme. Intact 28S and 18S rRNAs extracted from human cells (**Figure S.1D-E**), assembled large (LSU) (**Figure 3B**) and small (SSU) (**Figure S.1F**) ribosomal subunits, and polysomes (**Figure 3C**) all induce changes in the hemin Soret bands, which is indicative of heme-rRNA interactions. The combined data are consistent with a model in which rRNA tentacles of human ribosomes bind to hemin *in vitro*.

**Figure 3.**
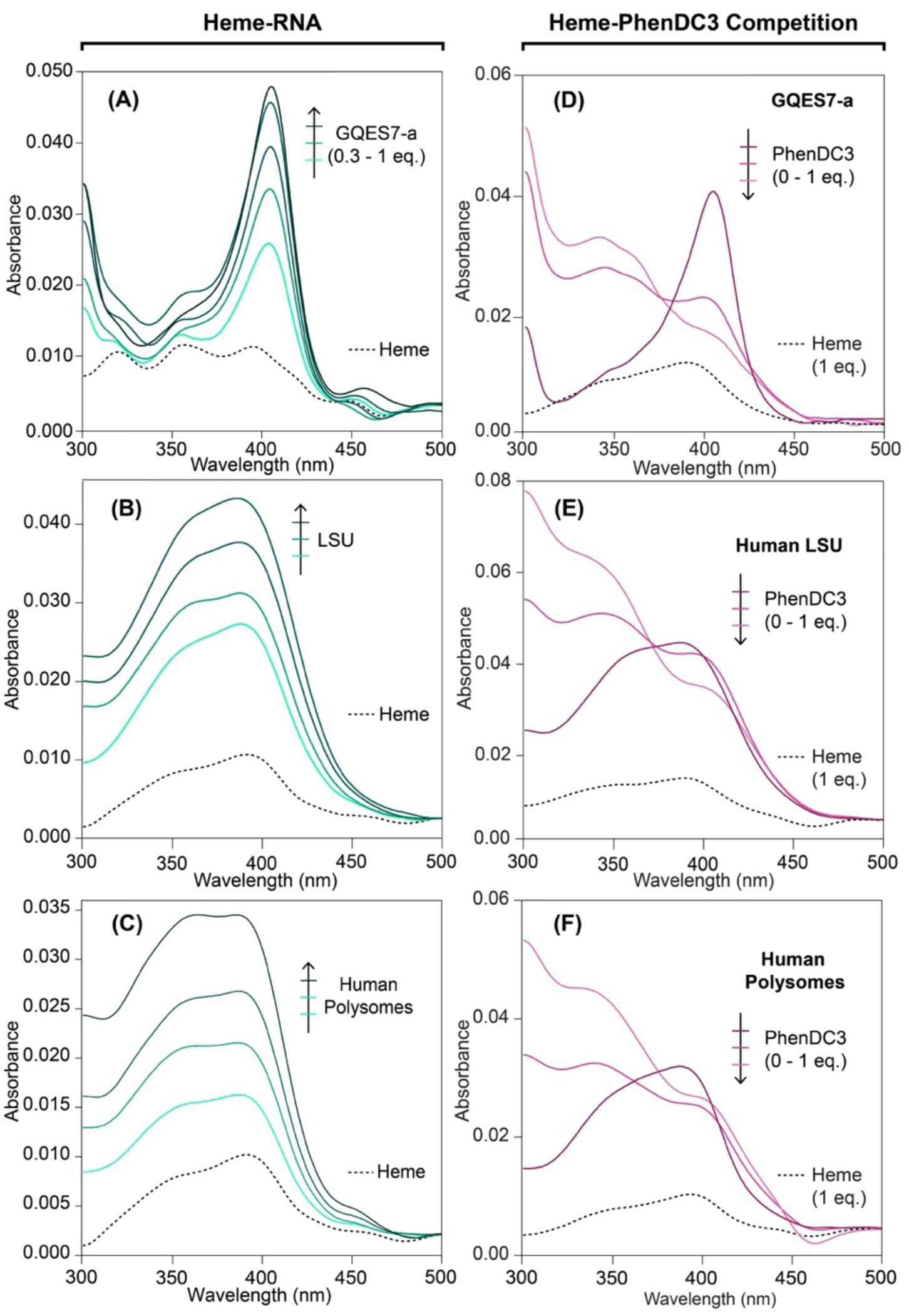
Human rRNA G4s bind heme *in vitro*. (A) UV-Vis spectra of heme during a titration with GQES7-a, (B) during a titration with the assembled LSU, and (C) during a titration with polysomes. (D) UV-Vis spectra of constant heme/GQES7-a during a titration with PhenDC3, (E) heme/LSU during a titration with PhenDC3, and (F) heme/polysomes during a titration with PhenDC3.

PhenDC3 was used to confirm binding of hemin to ribosomal G4s. PhenDC3, like hemin, end-stacks on G4s (29) and therefore competes with heme for binding to G4s. With fixed GQES7-a and hemin, addition of PhenDC3 causes a decrease in the intensity of the hemin Soret peak (**Figure 3D**) due to dissociation of heme. The same phenomenon is observed with assembled ribosomal particles (LSU: **Figure 3E**, SSU: **Figure S.2A**) and with polysomes (**Figure 3F**). Hemin that is associated with purified 28S and 18S rRNAs is also dissociated by PhenDC3 (**Figures S.2B-C**). Solutions of hemin with *mut*es3, however, do not show a change in the Soret peak upon the addition of PhenDC3 (**Figure S.2D**). Addition of PhenDC3 causes a slight increase in absorbance at 350 nm (**Figures 3D-F**). This phenomenon is intrinsic to PhenDC3, which absorbs at this wavelength (**Figure S.2E**). Taken together, the results here provide strong support for association of hemin with G4s on human ribosomes *in vitro*.

### Human ribosomes bind heme in vivo

We developed an assay that exploits differential interactions with hemin-agarose, an agarose resin covalently linked to heme, to report *in vivo* heme binding to ribosomes and rRNA. The degree to which any biomolecule interacts with heme in cells is inversely correlated with the extent to which it interacts with hemin-agarose upon lysis due to competition between endogenous heme and hemin-agarose for the heme-binding site. Therefore, the effects of heme binding factors *in vivo* can be monitored by determining if their interaction with hemin-agarose changes upon depletion of intracellular heme.

Accordingly, HEK293 cells were conditioned with and without succinylacetone (SA (33)), an inhibitor of heme biosynthesis. Lysates of these cells were incubated with hemin-agarose, and hemin-agarose interacting rRNA was quantified by RT-qPCR. Consistent with previous work (34), treatment with 0.5 mM SA for 24 hours caused a 7-fold decrease in total cellular heme in HEK293 cells (results not shown). The results reveal that rRNA binding to hemin-agarose relative to control agarose lacking heme increases by ∼4-fold in cells depleted of heme (**Figure 4A**). This result suggests that, under heme-depleted conditions, a greater fraction of rRNA heme binding sites are free and available to bind hemin agarose. In short, the data are consistent with a model in which ribosomal RNAs associate with endogenous heme.

**Figure 4.**
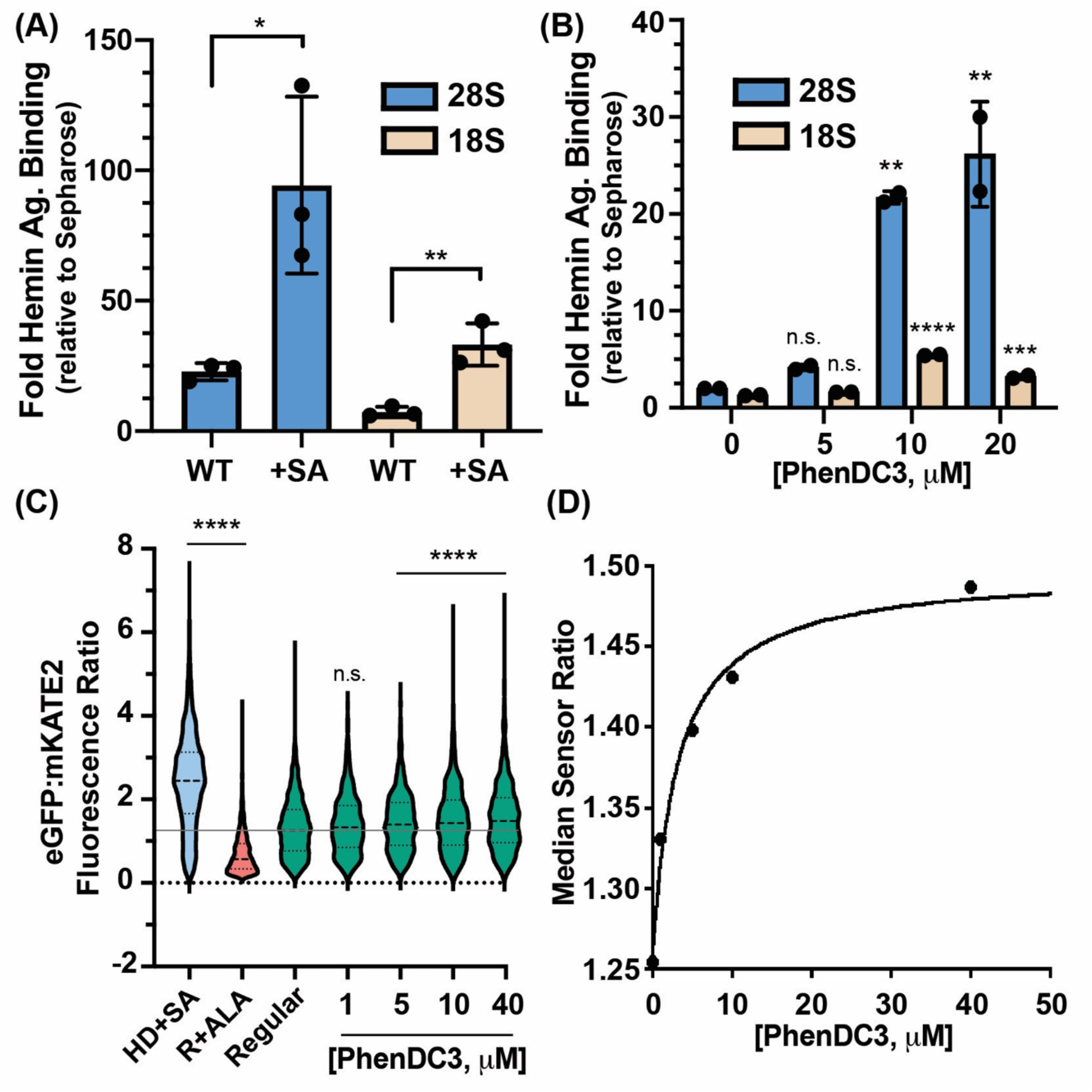
Ribosomes appropriate heme *in vivo* through rRNA G4s. (A) RT-qPCR analysis from untreated (WT) and SA-treated human cells. Statistical significance relative to WT is represented by asterisks using Student’s *t*-test. Each dot represents a biological replicate. (B) RT-qPCR analysis from PhenDC3-treated HEK293 cells. Statistical significance relative to no treatment conditions is represented by asterisks using ordinary one-way ANOVA with Dunnett’s post-hoc test. Each dot represents a technical replicate coming from individual biological replicates. The experiment was performed a total of 2 times with similar dose-dependent trends (**Fig. S.3A**). Data in (A) and (B) are represented as RNA enrichment in hemin agarose beads relative to control sepharose beads. (C) Single cell analysis of HS1-transfected HEK293 cells grown in heme deficient media containing succinylacetone (HD+SA), regular media containing 5-aminolevulinic acid (R +ALA), or regular media (regular) with the indicated concentrations of PhenDC3. Statistical significance relative to regular conditions is represented by asterisks using the Kruskal-Wallis ANOVA with Dunn’s post-hoc test. * P < 0.05; ** P < 0.01; *** P < 0.001; **** P < 0.0001; n.s. = not significant; (n ≅ 1500 cells). (D) Median HS1 sensor ratios obtained in (C) as a function to PhenDC3 concentration.

### In vivo PhenDC3 increases binding of ribosomes to hemin agarose

To determine if rRNA G4s bind heme *in vivo*, we treated HEK293 cells with the G4 ligand PhenDC3 (48 hrs at 37 °C). PhenDC3 and heme compete for binding to G4 rRNA *in vitro* (**Figure 3**). Thus, if rRNA G4s bind heme *in vivo*, PhenDC3 is expected to displace any rRNA bound heme. After cell lysis, excess hemin agarose is expected to out compete rRNA bound PhenDC3. Indeed, RT-qPCR reveals that PhenDC3 treatment of HEK293 cells causes a dose-dependent increase in binding of the LSU to hemin agarose (**Figure 4B**). A corresponding, but weaker signal is seen for the SSU, in agreement with the higher abundance of G4 regions in the LSU than in the SSU (**Figure 1A**). Treatment of HEK293 cells with carrier DMSO does not result in hemin agarose enrichment (**Figure S.3B**) or in heme-G4 interaction alterations (**Figure S.3C**). Taken together, these results indicate that G4s in rRNA bind heme in cells.

### rRNA G4s regulate heme bioavailability in vivo

To determine if rRNA G4s regulate heme homeostasis, we deployed a previously described genetically encoded ratiometric fluorescent <u>h</u>eme <u>s</u>ensor, HS1. HS1 is a tri-domain fusion protein consisting of a heme binding domain, cytochrome *b*_562_, fused to fluorescent proteins, eGFP and mKATE2, whose fluorescence is quenched or unaffected by heme, respectively. Thus, the eGFP:mKATE2 fluorescence ratio is inversely correlated with bioavailable heme as measured by HS1. HS1 was previously used to characterize heme homeostasis in yeast, bacteria, and mammalian cells, and was instrumental in identifying new heme trafficking factors and signals that alter heme biodistribution and dynamics (33, 35, 36). We asked if cytosolic heme bioavailablity is altered in response to PhenDC3 (33). As indicated in **Figure 4C**, single cell analysis of a population of ∼1500 HEK293 cells per condition indicate the median HS1 eGFP/mKATE2 ratio increases upon heme depletion in heme deficient media containing SA (HD+SA) and decreases upon increasing intracellular heme when cells are conditioned with the heme biosynthetic precursor 5-aminolevulinic acid (ALA) to drive heme synthesis. Titration of PhenDC3 results in a dose dependent increase in HS1 sensor ratio, indicating heme is less bioavailable when it is displaced from G4s in rRNA. The fractional heme saturation of HS1 decreases by ∼15% (**Figure 4D**). Together, our data indicate that rRNA G4s bind heme and regulate intracellular heme bioavailability.

## DISCUSSION

The results here provide strong evidence that ribosomal tentacles form G4s in human cells, and that these G4s are involved in appropriating heme. Immunofluorescence experiments with BG4 and L19 antibodies suggest a specialized fraction of cytosolic ribosomes (∼5%) form G4s and that most extra-nuclear G4s (∼83%) are ribosomal. The small fraction of ribosomes observed to form G4s *in vivo* contrasts with the high stability of ribosomal G4s *in vitro* (7, 8). This difference is consistent with Guo and Bartel, who propose that eukaryotic cells have a robust machinery that unfolds G4s (16). However, the high concentration of rRNA acts in opposition to the low frequency per ribosome, so the RNA G4s are reasonably abundant. The RNA G-quadruplexome appears to be ribosome-centric.

We previously reported that surfaces of both the SSU and the LSU contain G4 sequences (7, 8). A broad variety of data are consistent with more extensive formation of G4s on the LSU than on the SSU. These data include:

∘ more abundant and more expansive G-tracts on the LSU than the SSU (8),
∘ greater conservation over phylogeny of LSU G-tracts than SSU G-tracts (7, 8),
∘ higher thermodynamic stability of LSU G4s than SSU G4s (7, 8),
∘ greater heme binding to G4 oligomers from the LSU than those from the SSU (**Figure 3A, Figure S.1B**),
∘ greater enrichment of LSU than SSU particles in BioTASQ pulldowns (**Figure 2D**),
∘ greater enrichment of LSU than SSU particles in hemin-agarose pulldowns (**Figure 4A**), and
∘ greater effect of *in vivo* PhenDC3 treatment on LSU than on SSU rRNA in hemin-agarose pulldowns (**Figure 4B**).

Our findings that rRNA G4 forms complexes with heme *in vivo* has major implications for the physiology of G4-heme interactions. Decades of *in vitro* biophysical and chemical characterization of G4-heme complexes have found that they interact with high affinity (*K*_D_ ∼ 10 nM) and are potent redox catalysts, facilitating peroxidase and peroxygenase reactions (4-6). However, it remained unclear if heme-G4 complexes are formed *<u>in vivo</u>* and if heme-G4 catalyzed reactions were physiologically relevant.

Gray and coworkers (29) recently proposed that heme binds to G4s *in vivo*, based on the transcriptional response of cells to PhenDC3. PhenDC3 up-regulates heme degrading enzymes like heme oxygenase, and other iron and heme homeostatic factors. These responses were interpreted to support a model in which G4s sequester and detoxify heme in cells (29). Here, by exploiting differential interactions of *apo*-rRNA and heme-rRNA with hemin-agarose, we developed more direct methods to establish *in vivo* heme-G4 interactions, with a focus on rRNA G4s. Our results are consistent with the work of Gray *et. al.* in that we too conclude that G4s bind hemin *in vivo*. Moreover, our observation of rRNA-heme interactions *in vivo* supports a physiological role of rRNA G4-heme complexes in redox chemistry (37, 38). Indeed, work by Sen concurrent with ours has established G4-heme interactions *in vivo* by exploiting the peroxidase activity of G4-hemin complexes to self-biotinylate G4s in RNA and DNA using a phenolic-biotin derivative.

We propose that heme-rRNA G4 interactions may be important for protein hemylation reactions or buffering cytosolic heme. Indeed, PhenDC3, which competes for heme binding in G4s, causes a decrease in heme bioavailability as measured by the heme fluorescent sensor, HS1. This could be due to displacement of heme from rRNA G4s to a site that is less exchange labile, resulting in the observed decrease in HS1 heme binding. Alternatively, the upregulation in heme oxygenase due to PhenDC3 treatment (29) may decrease cellular heme levels, giving rise to the observed decrease in bioavailable heme. Overall, our results indicate rRNA G4s are sites of exchangeable heme in cells that may be available for heme dependent processes and hemylation reactions at the ribosome.

Taken together, the results here suggest that structural features of the human ribosome coupled with its high cytosolic abundance facilitate association with endogenous hemin. These results provide potentially new insights into the molecules and mechanisms underlying intracellular heme trafficking and bioavailability, which are currently poorly understood (1-3, 39). Our results suggest that ribosomes, and G4 containing rRNA in particular, are central hubs of heme metabolism, acting to buffer intracellular heme and possibly regulate heme trafficking and cotranslational hemylation. The ribosome as a hub for heme is consistent with its role as an abundant and versatile sink for ions and small molecules, including antibiotics, (40) platinum-based drugs, (41-43) certain metabolites (44), and metal cations Mg^2+^, Ca^2+^, Mn^2+^, and Fe^2+^, K^+^ (45-51).

## MATERIALS AND METHODS

### Cell culture

HEK293 cells were cultured in Dulbecco’s Modified Eagle Media (DMEM) containing 4.5 g/L Glucose without Sodium Pyruvate and L-Glutamine (Corning) supplemented with 10% fetal bovine serum (FBS) (Corning) and 2% penicillin-streptomycin solution (Gibco) in a humidified incubator kept at 37 °C with a 5% carbon dioxide atmosphere.

### RNAs

GQES7-a and GQES7-b were synthesized *in vitro* by transcription (HiScribe™ T7 High Yield RNA Synthesis Kit; New England Biolabs). GQes3 and *mut*es3, were purchased from Integrated DNA Technologies. Human 28S and 18S rRNAs were extracted from HEK293 cells with TRIzol (Invitrogen). Intact rRNAs were isolated by pipetting from a native agarose gel after running the rRNA into wells in the center of the gel. The rRNA was then precipitated with 5 M ammonium acetate-acetic acid (pH 7.5) with excess ethanol. RNA sequences are listed in Table S.1.

### RNA Annealing

RNAs were annealed by heating at 95°C for 5 min and cooled to 25°C at 1°C/min and incubated for 10 min at 4°C.

### UV-Visible Absorbance Heme-RNA Binding

Stock solutions of hemin chloride (1mM) were prepared in DMSO. Prior to use, the hemin chloride solution was sonicated for 10 min. RNAs (GQES7-a, GQES7-b, GQes3, *mut*es3) were annealed as described above in 50 mM KCl and 10 mM Tris-HCl, pH 7.5 in increasing RNA concentrations (for rRNA oligomers: from 0.3 to 1 equivalents of heme). The annealing buffer for intact 28S and 18S rRNAs and assembled ribosomal subunits and polysomes was the same as that of the rRNA oligomers except for the inclusion of 10 mM MgCl_2_. After binding, heme was added to a final concentration of 3 μM. Solution were allowed to stand at room temperature for 30 min then loaded onto a Corning® 384 Well Flat Clear Bottom Microplate. Absorbances were recorded from 300 nm to 700 nm on a a BioTek Synergy™ H4 Hybrid plate reader.

### UV-Visible Absorbance, Heme-PhenDC3 Competition Assay

For heme - PhenDC3 competition assaya, RNAs were annealed and allowed to bind to heme as above. Final heme concentration was 3 μM. Final RNA concentrations were: GQES7-a (3 μM), intact human 18S rRNA (65 nM), intact human 28S rRNA (22 nM). After solutions were inclubated for 30 min at room temperature, PhenDC3 or carrier DMSO was added to final concentrations consisting of 1.5 μM, 3 μM, and 6 μM. Samples were allowed to stand at room temperature for 15 minutes and were loaded onto a Corning® 384 Well Flat Clear Bottom Microplate. Absorbance was recorded from 300 nm to 700 nm.

### Total Heme Quantification of Untreated and SA-treated HEK293 cells

Heme was quantified as described. (52) Briefly, HEK293 cells were seeded in complete DMEM media at an initial confluency of 10% and incubated at 37 °C for 48 hrs. Media for SA-treated cells was replaced by DMEM supplemented with 10% heme-depleted FBS and 0.5 mM SA. Heme depletion of serum performed as described. (53) Media for untreated cells was replaced by complete media (supplemented with 10% regular FBS) and allowed to seed at 37 °C for 24 hrs. Cells were harvested by scrapping and counted using an automated TC10 cell counter (Bio-Rad). Then, 2.5×10^4^ cells per condition were treated with 20 mM oxalic acid and incubated at 4 °C overnight in the dark. An equal volume of 2 M oxalic acid was added to the cell suspensions. Samples were split, with half incubated at 95 °C for 30 min and half incubated at room temperature for 30 min. Samples were centrifuged at 21,000*g* for 2 min, and 200 μL of each was transferred to a black Greiner Bio-one flat bottom fluorescence plate, Porphyrin fluorescence (ex: 400 nm, em: 620 nm) was recorded on a Synergy Mx multi-modal plate reader. Heme concentration was calculated from a standard curve prepared by diluting a 0.1 μM hemin chloride stock solution in DMSO and treated as cell suspensions above. To calculate heme concentration, the fluorescence of the unboiled samples is taken as the background level of protoporphyrin IX and it is subtracted from the fluorescence of the boiled sample, which is used as the free base porphyrin produced upon the release of the heme iron. Using this method, our data suggest SA-treatment of HEK293 cells results in a 7-fold decrease in the total cellular heme concentration.

### Hemin Agarose Binding

HEK293 cells were seeded onto a 6-well plate at an initial confluency of 20% in Dulbecco’s modified Eagle’s medium (DMEM) with 10% Fetal Bovine Serum (FBS) and allowed to seed for 48 hrs at 37 °C. Media was then replaced for DMEM with 10% heme-depleted FBS supplemented with 0.5 mM succinyl acetone (for SA-treated cells). For untreated cells, media was changed for DMEM in 10% regular FBS. Both treated and untreated samples were allowed to incubate at 37 °C for 24 hrs. Cells were then collected by scrapping and lysed using 1.5 mm zirconium Beads (Benchmark). Lysates were quantified by Bradford assay. In the meantime, hemin agarose beads and sepharose beads were equilibrated 3 times by centrifugation with Lysis buffer (0.1% Triton X-100, 10 mM Sodium Phosphate, 50 mM KCl, 5 mM EDTA, pH 7.5, 1X protease arrest, RNasin RNase Inhibitor (Promega)). 100 μL of beads (50 μL bed volume) were used per biological replicate. After bead equilibration, each lysate was divided into two and 10 μg were loaded to hemin agarose and 10 μg to sepharose beads. Mixtures were allowed to bind for 60 min, rotating at 20 rpm at room temperature. Then, three washes were performed using Lysis buffer and supernatants were discarded. Each wash consisted of 10 min incubation at room temperature with 20 rpm rotation followed by centrifugation at 700*g* for 5 min. Bead bound fractions were eluted by a 15 min incubation at room temperature with 20 rpm rotation in 50 μL of 1M imidazole in Lysis buffer followed by centrifugation at max. speed for 2 min and supernatants were collected. RNA was then extracted from eluted fractions with TRIZOL using the manufacturer’s protocol. For the PhenDC3 titration in HEK293 cells experiment, the same protocol was followed with the difference that cells were allowed to seed for 24 hrs (20% initial confluency) and then PhenDC3 was added in increasing concentrations (5 μM, 10 μM, 20 μM). DMSO carrier treatment was performed the same way but with equivalent DMSO volumes. Cells were left at 37 °C for 48 hrs and collected and lysed as described above.

### RT-qPCR

The sets of primers used can be found in Table S.2. Luna Universal One-Step RT-qPCR kit (New England Biolabs) was used following the manufacturer’s protocol. Fold enrichments were calculated by comparing the C(t) values obtained from RNAs extracted from hemin agarose to RNAs extracted from sepharose beads. Three biological replicates were performed for all the RT-qPCR experiments. For BioTASQ experiments, fold enrichments were calculated by comparing the C(t) values obtained from the lysates containing BioTASQ + beads with those containing beads only.

### Heme Bioavailability Assay using the HS1 Sensor

HEK293 cells were plated and transfected in polystyrene coated sterile 6 well plates (Grenier) for flow cytometry. The cells were plated in basal growth medium Dulbecco’s modified eagle medium (DMEM) containing 10% fetal bovine serum. At 30% confluency cells were transfected with the heme sensor plasmid pEF52α-hHS1 using Lipofectamine LTX according to the manufacturer’s protocols. After 48 hours of treatment with transfection reagents, cells treated with PhenDC3 (1 mM stock) in fresh DMEM 10% FBS for 24 hours prior to harvesting. Heme depleted cells were treated with 500 μM succinylacetone (SA) in DMEM containing 10% heme depleted FBS for 72 hours prior to harvesting. Heme sufficient cells were treated with 350 μM 5-aminolevulinic acid (ALA) in DMEM 10% FBS for 24 hours. Cells were harvested in 1X PBS for flow analysis. Flow cytometric measurements were performed using a BD FACS Aria Ill Cell Sorter equipped with an argon laser (ex 488 nm) and yellow-green laser (ex 561 nm). EGFP was excited using the argon laser and was measured using a 530/30 nm bandpass filter, mKATE2 was excited using the yellow-green laser and was measured using a 610/20 nm bandpass filter. Data evaluation was conducted using FlowJo v10.4.2 software. Single cells used in the analysis were selected for by first gating for forward scatter (FSC) and side scatter (SSC), consistent with intact cells, and then by gating for cells with mKATE2 fluorescence intensities above background were selected. The fraction of sensor bound to heme may be quantified according to the following equation (33):

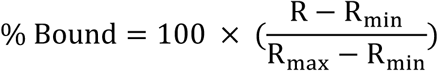

where R is the median eGFP/mKATE2 fluorescence ratio in regular media and *R*_min_ and *R*_max_ are the median sensor ratios when the sensor is depleted of heme or saturated with heme. R_min_ and R_max_ values are derived from cells cultured in heme deficient media conditioned with succinylacetone (HD+SA) or in media conditioned with ALA (33). The plot in Figure 4D was obtained by fitting the median sensor ratios in Figure 4C to the following 1-site binding model (33, 54):

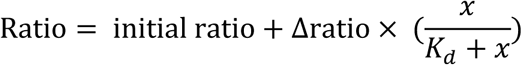

where x is the independent variable, [PhenDC3]

### BG4 purification

pSANG10-3F-BG4 was a gift from Shankar Balasubramanian (Addgene plasmid # 55756; http://n2t.net/addgene:55756; RRID:Addgene_55756). BL21 cells transformed with this plasmid were grown in room temperature and induced overnight with 0.1mM IPTG. Cells were pelleted, then resuspended in xTtractor (Takara) supplemented with Protease arrest (G-protein), lysozyme and DNase I. Sonicated cell lysate was combined with Ni-NTA resin (Invitrogen) and purified via the his-tag. BG4 was further purified by FPLC using a Superdex75 size exclusion column (GE Healthcare).

### Immunofluorescence

Immunofluorescence was performed by standard protocols. HEK293 cells were seeded onto Poly-L-lysine coated cover glass two days before the experiment and fixed in 4% formaldehyde for 15 min. Cells were permeabilized with 0.1% Triton X-100 for 3 min and blocked with 5% donkey serum (Jackson ImmunoResearch), followed by incubation with antibodies for 1 hr at room temperature or overnight at 4 °C. Antibodies used here are: BG4, rabbit anti-FLAG (Cell Signaling Technology, 14793S), mouse anti-L19 (Santa Cruz Biotechnology, sc-100830), mouse anti-rRNA (Santa Cruz Biotechnology, sc-33678), mouse anti-Calnexin (Santa Cruz Biotechnology, sc-23954), Alexa Fluor 488 conjugated donkey anti-rabbit (Jackson ImmunoResearch, 711-545-152), Rhodamine Red-X conjugated donkey anti-mouse (Jackson ImmunoResearch, 715-295150). After staining cells were carefully washed with DPBS supplemented with 0.1% tween-20. Nuclear DNA was stained with 4’,6-diamidino-2-phenylindole (DAPI). Images were acquired with a Zeiss 700 Laser Scanning Confocal Microscope. PhenDC3 treatment consisted of incubation at 37 °C overnight at 10 μM final PhenDC3 concentration prior to cell fixation. Determination of colocalization ratios was performed as described in Zen software (Zeiss). No primary antibody controls as well as RNase A and PhenDC3 treated images are reported in **Figures S.4** and **S.5**. The “Colocalization” image in **Figure 2A,B** is showing the G4 signal that colocalizes with L19 and with the ER (yellow pixels) and the one that does not colocalize (green pixels). “L19”, “ER”, and “BG4” images only present their respective fluorescence signals.

### BioTASQ capture of cellular RNAs

BioTASQ experiments followed published protocols *in vitro* (8) and *in vivo* (18). Briefly, HEK293 cells were seeded onto a 6-well plate at 20% confluency and allowed to incubate at 37 °C for 48 hrs. Cells were then crosslinked with 1% paraformaldehyde/PBS for 5 min at room temperature. Crosslinking was stopped by incubating cells with 0.125 M glycine for 5 min at room temperature. Cells were harvested by scrapping and resuspended in Lysis Buffer (200 mM KCl, 25 mM Tris-HCl, pH 7.5, 5 mM EDTA, 0.5 mM DTT, 1% Triton X-100, RNasin RNAse Inhibitor, 1X protease arrest). Cells were lysed by sonication (30% amplitude, 10 sec. on and off intervals, 2 min sonication time). The lysate was then split: BioTASQ was added at a final concentration of 100 μM to one of the samples, the other one was left untreated. Lysates were incubated at 4 °C overnight with gentle rotation. Sera-Mag magnetic streptavidin-coated beads (GE Healthcare) were washed three times with wash buffer (5 mM Tris-HCl pH 7.5, 0.5 mM EDTA, 1 M KCl). Each wash was followed by centrifugation at 3,500 rpm for 5 min at 4 °C. Beads were then treated with Buffer 1 (0.1 M NaOH, 0.05 M KCl in RNase/DNase-free water) two times at room temperature for 2 min and then centrifuged at 3,500 rpm at 4 °C, 5 min, and washed with Buffer 2 (0.1 M KCl in RNase/DNase-free water). Lastly, to block, beads were treated with 1 μg/mL BSA and 1 μg/mL yeast tRNA and allowed to incubate at 4 °C overnight with gentle rotation.

After incubation overnight with BioTASQ, cell lysates were treated with 1% BSA for 1 hr at 4 °C. Washed magnetic beads were added to the lysates (20 μg beads /sample) and allowed to mix with gentle rotation at 4 °C for 1 hr. Beads were then washed three times with Lysis buffer for 5 min and then crosslinking was reversed by incubating the beads at 70 °C for 1 hr. Finally, TRIZOL was used to extract RNAs, for analysis by RT-qPCR.

## ACKNOWLEDGMENTS

The authors thank Drs. Rebecca Donegan, Jonathan B. Chaires, David Monchaud and Judy Wong, and Claudia Montllor-Albalate for helpful discussions. We acknowledge Andrew Shaw and the core facilities at the Parker H. Petit Institute for Bioengineering and Bioscience at the Georgia Institute of Technology for expert advice and the use of equipment. Purified human ribosomes and polysomes were a gift from Immagina BioTechnology. BioTASQ was a gift from Dr. David Monchaud. This work was supported by NASA, 80NSSC17K0295 and 80NSSC18K1139 (Center for the Origin of Life) to LDW, the NIH, ES025661 to ARR, and the NSF, MCB-1552791 to ARR.

## AUTHOR CONTRIBUTIONS

SMF, CI, CMM, ARR and LDW conceived and designed the experiments; SMF, CI and CMM performed the experiments; SMF, CI, CMM, ARR, and LDW analyzed data; SMF, ARR and LDW prepared figures; and SMF, ARR and LDW wrote the paper.

## SUPPLEMENTARY MATERIALS

**Figure S.1.**
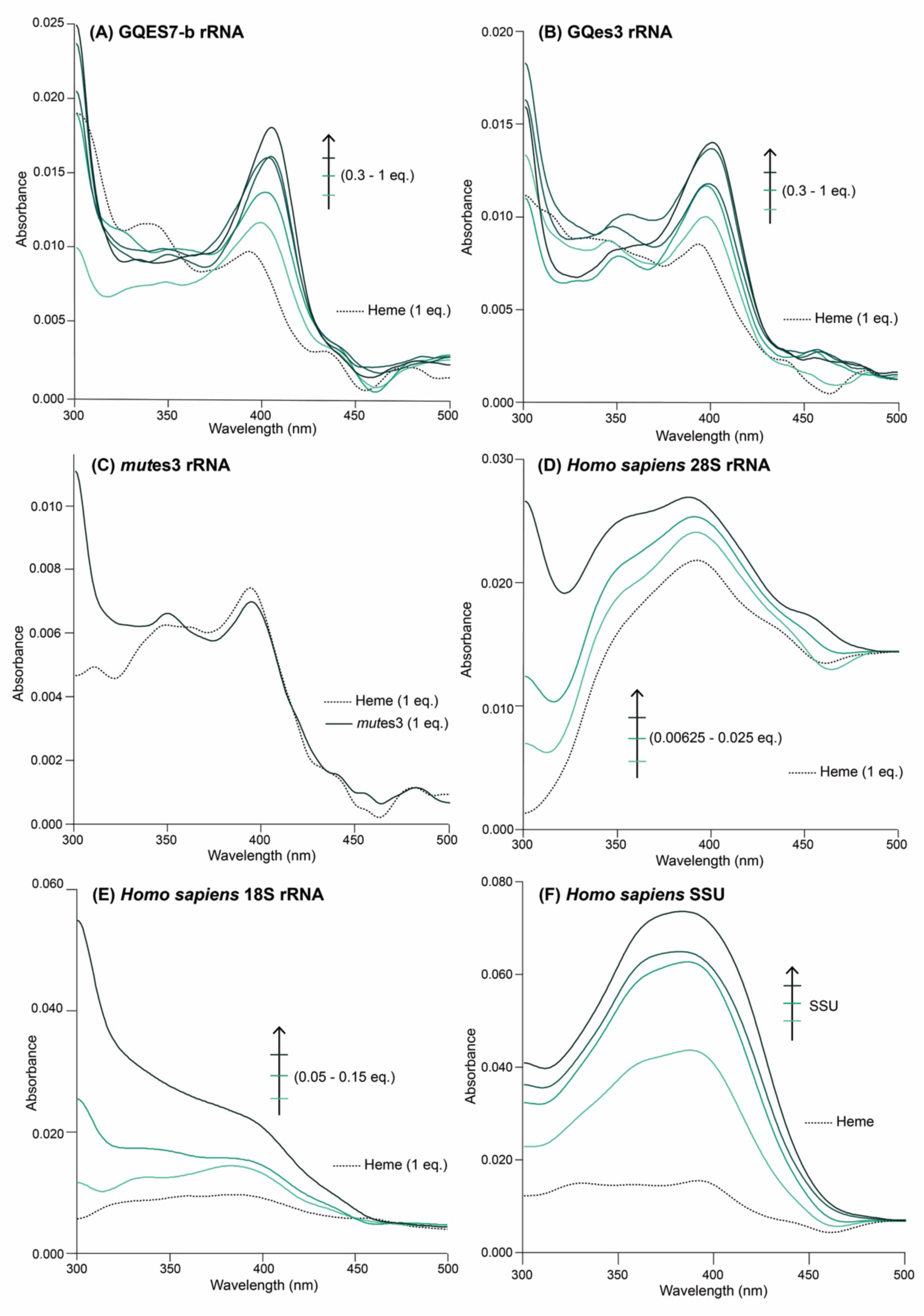
UV-Vis spectra of increasing concentrations of GQES7-b (A), GQes3 (B), *mut*es3 (C), intact human 28S rRNA (D), intact human 18S rRNA (E), and assembled small subunit (F) to a constant concentration of heme.

**Figure S.2.**
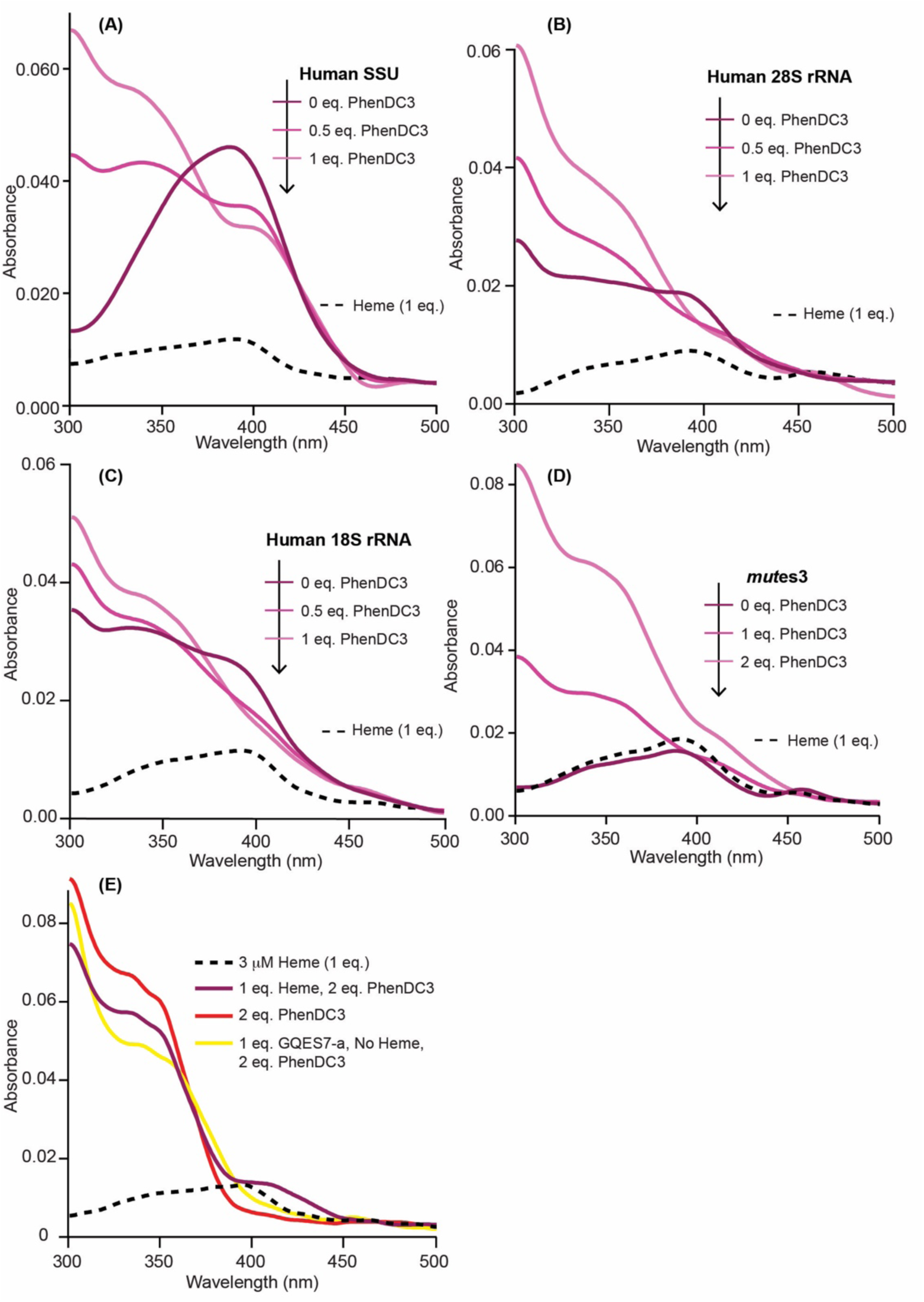
UV-Vis spectra of titration of PhenDC3 to constant heme and (A) human assembled SSU, (B) intact human 28S rRNA, (C) intact human 18S rRNA, and (D) and *mut*es3. (E) UV-Vis spectra of PhenDC3 reveals peak at 350 nm is intrinsic of PhenDC3.

**Figure S.3.**
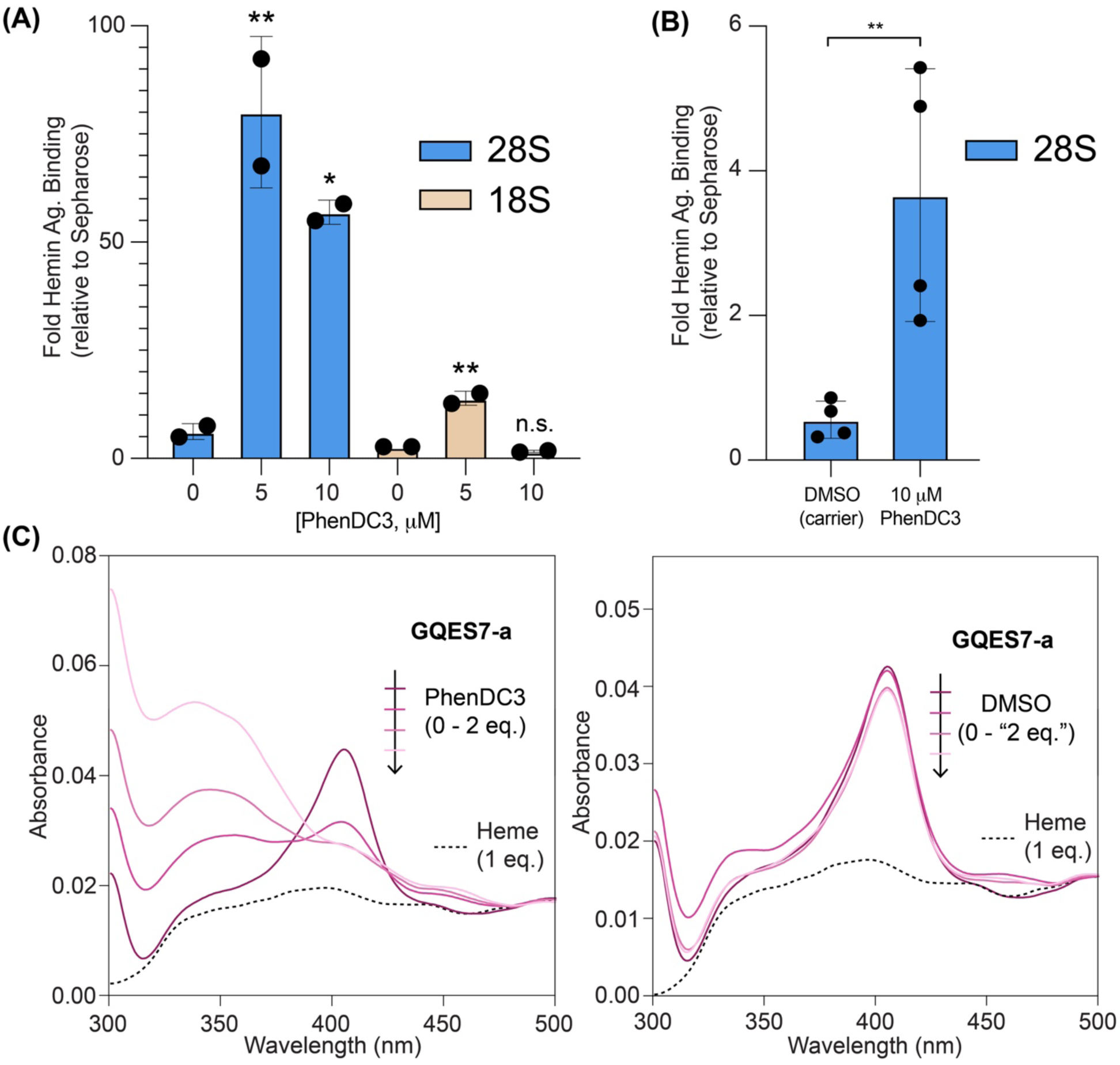
(A) RT-qPCR analysis oif 28S and 18S rRNAs from PhenDC3-treated HEK293 cells. PhenDC3 treatment consisted of 48 hrs at 37 °C in the concentrations listed in the figure. Each dot represents a technical replicate coming from individual biological replicates. (B) RT-qPCR analysis of 28S rRNA from HEK293 cells treated with 10 μM PhenDC3 or carrier DMSO. Each dot in (B) represents a technical replicate coming from 2 biological replicates. (C) UV-Vis spectra of increasing concentrations of PhenDC3 or carrier DMSO to constant heme and GQES7-a show DMSO does not affect the binding of heme to ribosomal G4s. Data in (A) and (B) are represented as RNA enrichment in hemin agarose beads relative to control sepharose beads. Statistical significance is represented by asterisks using ordinary one-way ANOVA with Dunnett’s post-hoc test. * P < 0.05; ** P < 0.01.

**Figure S.4.**
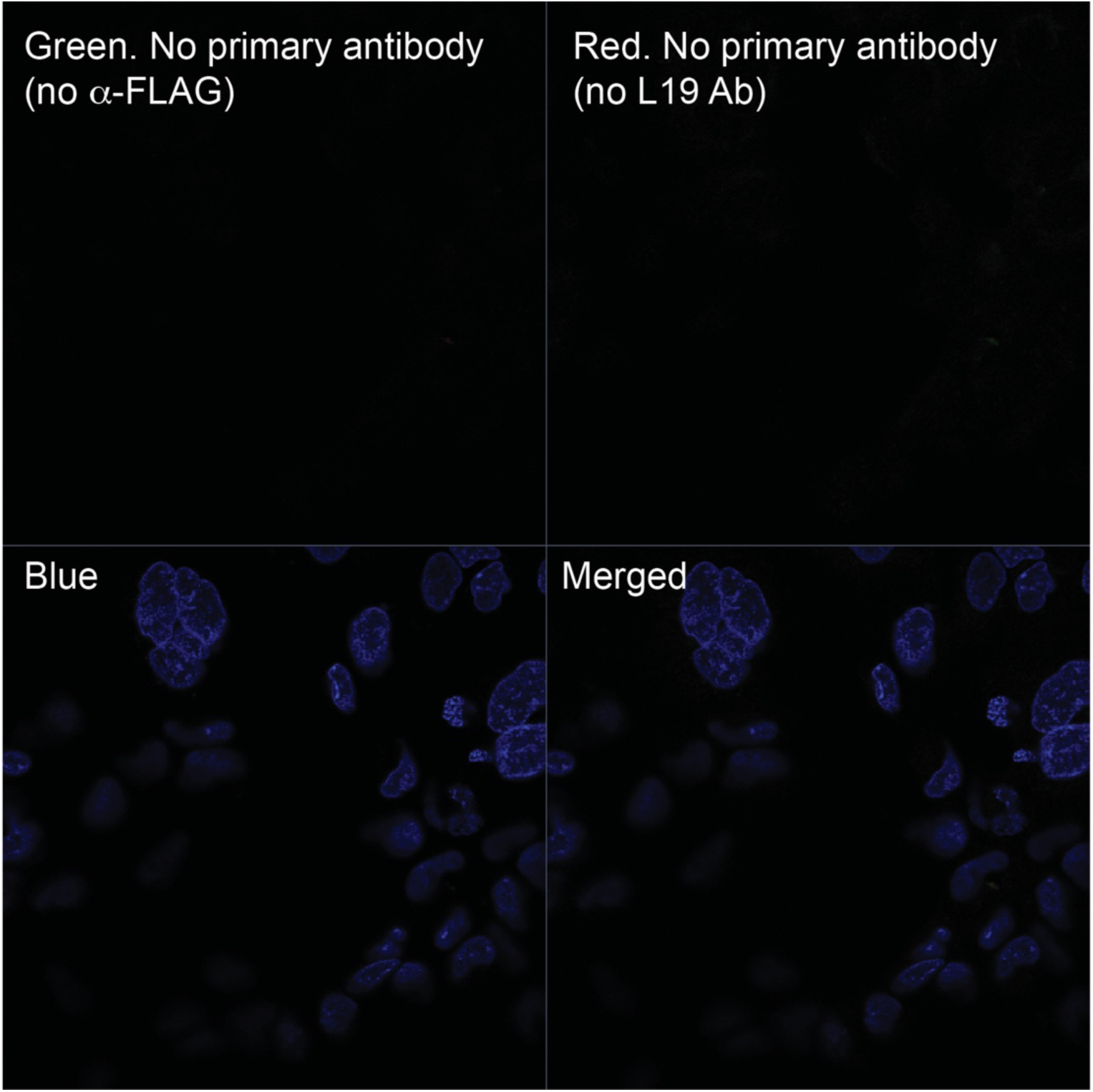
Confocal microscopy images of fixed HEK293 cells without the primary antibodies *α*-FLAG (green channel) and without *α*-L19 antibody (red channel). Results demonstrate signals obtained in confocal microscopy images are coming from the primary antibodies.

**Figure S.5.**
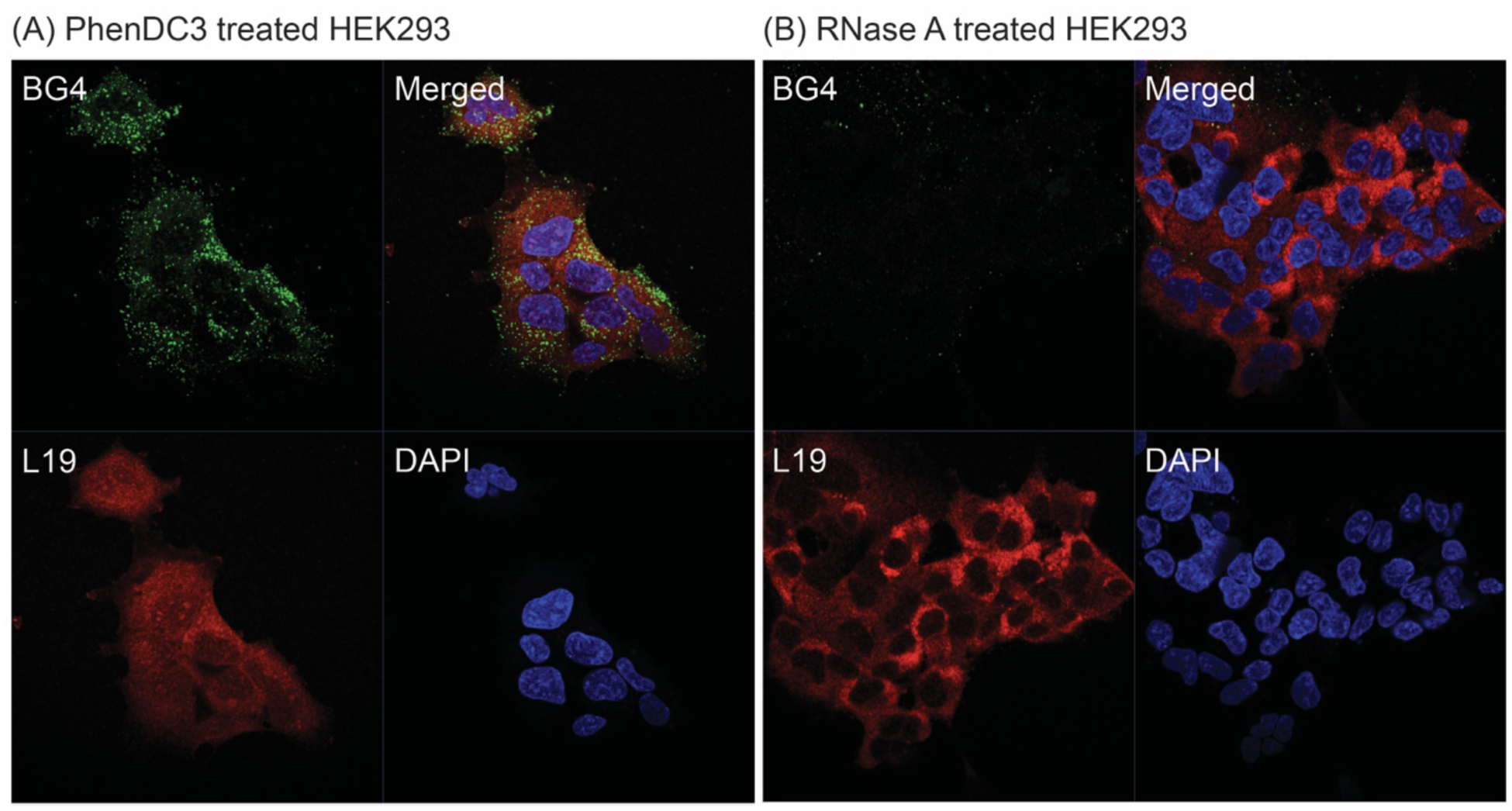
Confocal microscopy images of fixed HEK293 cells treated with (A) PhenDC3 or (B) RNase A. Results demonstrate BG4 signal is coming from cellular RNA G-quadruplexes. Note that “merged” images presented here are not shown in terms of BG4-L19 colocalization.

**Table S.1.**
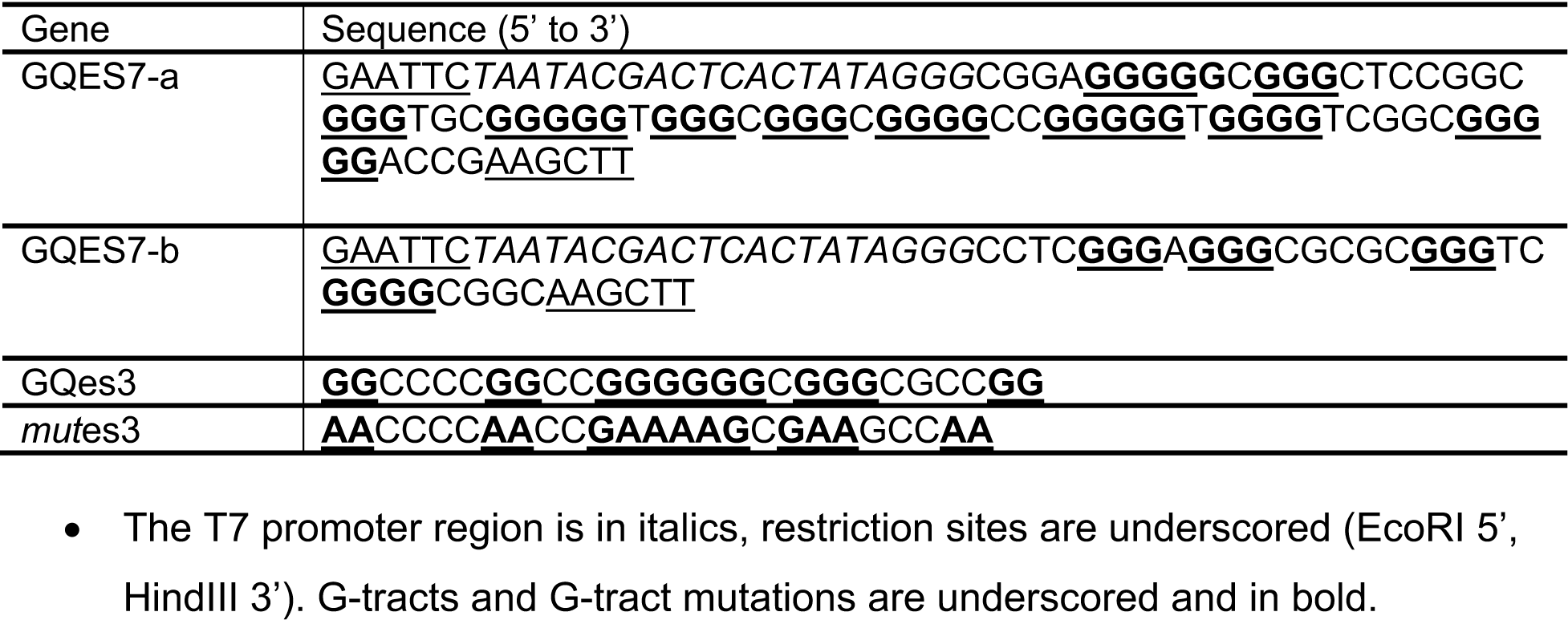
DNA and RNA sequences encoding RNAs used.

**Table S.2.**
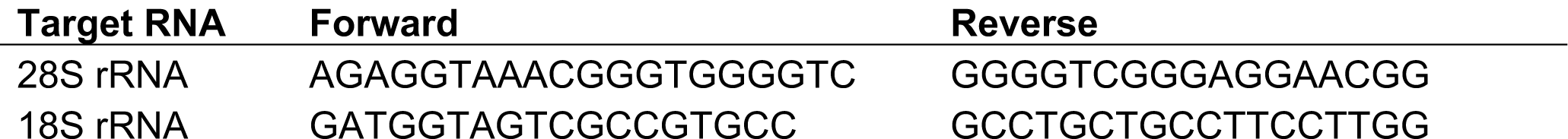
Primer sets used for RT-qPCR

